# Rhesus macaques model human Mayaro virus disease and transmit to *Aedes aegypti* mosquitoes

**DOI:** 10.1101/2025.04.16.649064

**Authors:** Adam J. Moore, Koen K. A. Van Rompay, William Louie, Jennifer K. Watanabe, Sunny An, Rochelle Leung, Jodie L. Usachenko, Peter N. Chu, Katherine J. Olstad, Colleen S. McCoy, Rafael K. Campos, Scott C. Weaver, Shannan L. Rossi, Lark L. Coffey

## Abstract

**Background:** Mayaro virus (MAYV) is a mosquito-borne alphavirus endemic to Latin America that causes fever and arthritis. Unlike the related chikungunya virus, MAYV has not caused widespread, human-amplified epidemics. One possible explanation is that human viremia levels are too low to support transmission to urban *Aedes* (*Stegomyia*) *aegypti* mosquitoes. We used rhesus macaques (RM) to model human-to-*Ae. aegypti* transmission and to further expand understanding of their relevance to human MAYV disease.

**Methodology/Principal Findings:** Twelve RM were inoculated with a genotype D lineage MAYV strain using one of 3 doses: 7 log_10_ plaque forming units (PFU) intravenously (IV), 7 log_10_ PFU subcutaneously (SC), or 3 log_10_ PFU SC. Viremia was measured daily in plasma and RM were euthanized 10- or 12-days post-inoculation (dpi). On 2, 3, 5, and 7 dpi, *Ae. aegypti* were allowed to bloodfeed, incubated for 10 days, then dissected and tested to detect MAYV in tissues and saliva. RM developed infectious MAYV viremias lasting 3 days, peaking 1-2 dpi with titers ranging from 2-6 log_10_ PFU/ml. RM inoculated with 7 log_10_ PFU IV developed significantly higher viremias (area under the curve) than those receiving 3 log_10_ PFU SC. MAYV RNA was detected in muscle, lymphoid, central nervous, and cardiac tissues. RM showed no signs of fever or joint swelling but some developed mild rashes in areas distant from mosquito feeding sites and histologic inflammation was observed in joints and muscles. Only *Ae. aegypti* that fed on viremic RM 2 dpi became infected, with an overall infection rate of 48%. Among all mosquitoes that fed on RM 2 dpi, only 2% (4/217) had infectious MAYV in their saliva, suggesting transmission competence. Despite 11 of 12 RM transmitting MAYV to at least one mosquito, individual RM varied in infectiousness to *Ae. aegypti,* and mosquito cohort infection rates did not correlate with RM viremia levels.

**Conclusions/Significance:** RM exhibit short-lived MAYV viremias, broad tissue tropism, and mild joint and muscle inflammation, closely resembling human infection. While viremic RM can infect *Ae. aegypti*, the transmission window is narrow and transmission by *Ae. aegypti* is rare. The combination of a short infectious period in RM and low transmissibility of *Ae. aegypti* infected from RM may help explain the absence of widespread urban MAYV outbreaks.

**AUTHOR SUMMARY:** Mayaro virus (MAYV) is a mosquito-borne virus found in Latin America that causes fever and joint pain, similar to chikungunya virus (CHIKV). However, unlike CHIKV, MAYV has not led to large outbreaks. One reason may be that levels of MAYV in human blood are too low for *Aedes aegypti* mosquitoes—known for spreading chikungunya, dengue, and Zika—to pick up and transmit the virus in cities. To better understand this, we studied rhesus macaques, a monkey species that serves as a model for how MAYV behaves in people. We tracked virus levels in their blood and tissues and observed mild joint and muscle inflammation, similar to Mayaro fever in people. Although the macaques were able to infect some *Ae. aegypti* mosquitoes, the transmission window was short, and only a few mosquitoes that became infected had virus in their saliva, suggesting transmission competence. This limited ability of urban mosquitoes to spread MAYV may help explain why major outbreaks have not occurred.

## INTRODUCTION

Mayaro virus (MAYV, *Togaviridae*, *Alphavirus;* species *Alphavirus mayaro*) is a mosquito-borne RNA virus that causes Mayaro fever, an acute febrile illness associated with chronic arthralgia [1]. Unlike the related chikungunya virus (CHIKV), MAYV has not caused widespread, human amplified, urban outbreaks in its enzootic regions of Latin America [1–3]. Consequently, research on MAYV has been relatively limited, especially compared to CHIKV. For example, although multiple vaccine candidates have been developed [4–9], none have progressed to evaluation in non-human primates (NHP). This is partly due to the historical lack of NHP models for MAYV. While NHP susceptibility was first reported in 1967 [10], it was not until 2023 that cynomolgus [11] and rhesus macaques (RM) [12] were developed as models of human MAYV disease. The RM study defined clinical disease, viremia kinetics, tissue tropism, and immune responses but was limited to three male animals inoculated with a single MAYV dose and employed a genotype L strain (where MAYV comprises 3 genotypes: D, L, and N), which has its known distribution restricted to Brazil [13] and Haiti [14]. Since disease outcomes can be influenced by dose, inoculation route, and host sex, one aim of this study was to identify MAYV outcomes in male and female subjects. If MAYV can efficiently be transmitted from viremic people to urban mosquitoes (human amplification), it poses a risk of major urban outbreaks. However, human MAYV viremia kinetics remain poorly characterized, making it unclear whether viremia levels are sufficient for transmission to urban mosquitoes. To address these gaps, we inoculated male and female RM once with one of 2 doses of a genotype D MAYV strain, which circulates throughout South America [13]. RM were inoculated subcutaneously to mimic infection from a feeding mosquito [15,16], or intravenously to bypass the skin and ensure infection, since, at the time we began these studies, neither of the 2023 papers demonstrating successful subcutaneous MAYV infection of macaques had been published. We then characterized RM viremia magnitude and duration, tissue tropism, neutralizing antibody levels, and histopathologic changes in muscles and joints.

The global spread of CHIKV is driven by transmission using *Aedes* (*Stegomyia*) *aegypti* and *Ae*. (*Stegomyia*) *albopictus* mosquitoes, which thrive in urban tropical environments and are expanding into temperate regions [17]. In contrast, MAYV outbreaks tend to be more localized [2,18], and seropositivity is higher among individuals living in or near forests than in cities [18–20]. This pattern suggests that human exposure primarily occurs in forested areas where MAYV cycles between non-human primates, small mammals, birds, reptiles, and multiple species of forest-dwelling mosquitoes [1,21,22]. Although a lack of urban mosquito surveillance in Latin America may obscure potential MAYV transmission in cities, some evidence suggests *Ae. aegypti* could play a role in MAYV transmission cycles. MAYV RNA has been detected in adult *Ae. aegypti* in Brazil [23,24], and infectious MAYV has been recovered from *Ae. aegypti* eggs [25]. Laboratory vector competence studies using artificial bloodmeals have also demonstrated *Ae. aegypti* are susceptible to MAYV [26–30]. Furthermore, *Ae. aegypti* that ingested MAYV in artificial bloodmeals developed detectable MAYV RNA in salivary glands and, in transmission experiments, cohorts of 10-15 mosquitoes that fed together on naïve immunocompromised mice 7 days later all transmitted MAYV to the mice, with 50% of naïve *Ae. aegypti* that subsequently fed on viremic mice becoming infected, demonstrating successful *Ae. aegypti* -mouse- *Ae. aegypti* MAYV cycling [26]. These findings suggest that *Ae. aegypti* could contribute to urban MAYV transmission, although immunocompromised mice are less representative of human infection than RM. If MAYV can be efficiently transmitted from infected humans to *Ae. aegypti*, it poses a risk for major urban outbreaks as seen with CHIKV. Therefore, another goal of this study was to use experimentally infected RM to evaluate MAYV transmission to *Ae. aegypti*.

## METHODS

### Animal source, care, use, and sample collection

This study used 12 healthy adult Indian origin rhesus macaques (RM, *Macaca mulatta*), including nine males and three females, aged 3.7 to 6.6 years. All animals were from a type D retrovirus-, simian immunodeficiency virus-, and simian lymphocyte tropic virus type 1-free colony at the California National Primate Research Center (CNPRC). All animals were previously exposed to the flavivirus dengue virus, which is not known to impact MAYV infection outcomes. All RM passed a physical exam before study enrollment. Although the alphavirus western equine encephalitis virus has historically circulated in California, no environmental activity in California mosquito pools has been detected since 2007, despite ongoing annual surveillance [31]. RM were born and raised at CNPRC and had been housed indoors for over a year before study enrollment. All experimental procedures were conducted at CNPRC, which is accredited by the Association for Assessment and Accreditation of Laboratory Animal Care International (AAALAC). Animal care followed the guidelines outlined in the 2011 *Guide for the Care and Use of Laboratory Animals* by the Institute for Laboratory Animal Research. RM were housed indoors in stainless steel cages (Lab Products Inc., Seaford, DE), with cage sizes scaled to each animal according to national standards. Housing conditions included a 12-hour (h) light/dark cycle, temperatures of 64 to 84°F, and room humidity of 30 to 70%. Animals that were previously bonded were housed together. They had free access to water and were fed a commercial high-protein diet (Ralston Purina Co., St. Louis, MO) supplemented with fresh produce. The study was approved by the Institutional Animal Care and Use Committee of the University of California, Davis (Protocol number 23106). Animals were monitored daily for clinical signs, including behavior changes, rash or redness, rectal temperature, and joint swelling at time of sedation. For MAYV inoculation, blood collections, and mosquito feedings, RM were immobilized with ketamine HCl (Dechra Veterinary Products, Overland Park, KS) at 10 mg/kg that was injected intramuscularly in the left leg after overnight fasting. Blood samples were collected using venipuncture. A portion of ethylenediaminetetraacetic acid (EDTA)-anticoagulated whole blood was used for complete blood counts, performed by the CNPRC Clinical Laboratories. The remaining blood was processed by centrifugation at 900 x g for 10 minutes (m) to separate plasma from cells. Plasma was then centrifuged again at 900 x g for 10 m to remove residual cells, after which aliquots were immediately frozen at −80°C. Blood collected without anticoagulant was processed via centrifugation at 900 x g for 10 m to obtain serum. Cerebrospinal fluid was collected from the cervical region. Half of the RM in each cohort were selected for euthanasia on 10 or 12 dpi. Euthanasia was performed with an overdose of pentobarbital, followed by necropsy and tissue collection. Samples were collected from all major organ systems, with a focus on lymphoid tissues, joints, and muscles. A skin biopsy, referred to as the “skin inoculation site” was collected from the middle back region, corresponding to the location where the SC-inoculated animals were injected with MAYV. Tissues were processed using three methods: (i) snap-frozen in liquid nitrogen and stored at −80°C; (ii) preserved in RNALater (ThermoFisher, Waltham, MA) followed by incubation overnight at 4°C then transfer to −80°C for long-term storage, and (iii) fixed in 4% paraformaldehyde.

### MAYV source and inoculations

All 12 RM were inoculated with MAYV strain IQT4235. The virus was derived from an infectious clone plasmid created from the sequence of strain IQT4235, a genotype D lineage virus which predominates in Latin America [6]. The plasmid was derived from the sequence of a MAYV strain that was originally isolated from the serum of a febrile human in Iquitos, Peru in 1997 (GenBank: MK070491) [27]. Before *in vitro* transcription, the complete MAYV cDNA sequence in the plasmid was verified (Plasmidsaurus, Eugene, OR). The sequence in the plasmid was identical to the GenBank sequence except for 2 non-coding mutations (T6298A and G10797A). To generate infectious MAYV, the cDNA clone was linearized using the *PacI* restriction endonuclease (New England Biolabs, Ipswich, MA), transcribed *in vitro* into mRNA using the mMessage mMachine SP6 transcription kit (ThermoFisher, Waltham, MA), and electroporated into Vero cells using a BioRad Gene Pulser XCell (Hercules, CA) with the following settings: 125 Volts, 10 millisecond pulse length, 1 second intervals, 3 pulses. After *in vitro* transcription, the RNA was deep sequenced using the methodologies described below. Infectious MAYV was harvested from Vero cell culture supernatant 2 days post-electroporation, titrated, and stored at −70°C until use. To evaluate effects of dose and inoculation route, male RM were randomly divided into 3 groups. Since only 3 females were available, 1 female was assigned to each group. Each group received a single MAYV inoculation on day 0 with one of the following: 7 log_10_ IV, 7 log_10_ PFU SC, or 3 log_10_ PFU SC. The 7 log_10_ dose exceeds mosquito-delivered doses and was intended to ensure infection. The 3 log_10_ dose was intended to simulate a mosquito-delivered dose, where our prior studies with CHIKV [32] and studies by us [27] and others [28,29] for MAYV show that *Ae. aegypti* salivate between 1.0-2.5 log_10_ PFU or 1.0-4.0 log_10_ genomic RNAs into capillary tubes. IV injections were administered in the saphenous vein, while SC injections were administered in the upper back between the shoulder blades. Inocula were back-titrated by plaque assay to verify the administered dose. Blood samples were collected daily from 1-7 days post-inoculation (dpi) and again at euthanasia, 10 or 12 dpi.

### MAYV RNA isolation from macaque plasma and tissues

Frozen plasma and tissues were thawed at room temperature. For tissue processing, each sample was individually weighed, then combined with 500 µl Dulbecco’s modified eagle medium (DMEM) (GenClone, Waltham, MA) and a 4 mm glass bead (ThermoFisher, Waltham, MA).

Homogenization was performed using a Retsch MM400 Tissue Lyser (Haan, Germany) at 30 shakes/s for 5 min, followed by a 10 min rest. Samples were then rotated and homogenized again using the same conditions. To clarify the supernatant, homogenized tissues were centrifuged at 3000 g for 2 min. Thawed plasma or supernatant from tissues was used immediately after homogenization for plaque assays or RNA extractions.

### Tissue processing and histopathologic analyses

RM tissues were fixed in 4% paraformaldehyde and then paraffin embedded, sectioned, and stained with hematoxylin and eosin. A pathologist, blinded to treatment, evaluated joints, tendons, and muscles for pathologic changes using a quantitative scoring metric (**Supplemental Table 1**). Tissues from age- and sex-matched colony control animals not part of this study that were not MAYV exposed served as comparators.

### Mosquito source, presentation to rhesus macaques, sampling, and processing

Adult *Ae. aegypti* mosquitoes (aged 3-9 days) from the 33^rd^ generation of a colony originally collected in Los Angeles, CA, and identified morphologically in the field [33], were used in this study. Mosquitoes were reared in an insectary under controlled conditions (28°C, 75% humidity, 12hr light/dark cycle) with free access to 10% sucrose in water. All mosquitoes presented to RM were from the same cohort. Twenty-four hours prior to presentation, sucrose was replaced with sterile water. Mosquitoes were housed in pint sized containers and transported inside plastic shoeboxes from a rearing insectary to the CNPRC animal rooms. On 2, 3, 5, and 7 days post inoculation (dpi) of the RM, cohorts of ∼100 mixed-sex *Ae. aegypti* were presented to the shaved abdomens of anesthetized RM and allowed to feed for 10-15 min. Feeding was confirmed by visual observation of red abdomens. After feeding, mosquitoes were transported in shoeboxes placed inside sealed bags into an arthropod containment level 3 (ACL-3) facility.

Mosquitoes were anesthetized with CO_2_ and bloodfed females were separated from males and non-fed females on a cold table (Bioquip, Compton, CA). Bloodfed females were incubated at 28°C with 75% humidity with 12-h light/dark cycle for 10 days post feed (dpf), with continuous access to 10% sucrose. Nine to 20 mosquitoes per RM per feeding day were incubated. At 10 dpf, mosquitoes were anesthetized with CO_2_ for 10 s, immobilized on a chill table, and their legs and wings were removed. Saliva was collected using expectoration assays, where the proboscis was inserted into a capillary tube containing DMEM with 5% heat-inactivated fetal bovine serum (FBS) (56°C, 30 min) (GenClone, Waltham, MA) and 1% penicillin-streptomycin (P/S) (Genesee Scientific, El Cajon, CA) for 30 min to stimulate salivation. Mosquito tissues were placed in 500 µl DMEM supplemented with 5% FBS and 1% P/S, then homogenized in a TissueLyser II (Qiagen, Hilden, Germany) at 20 shakes/s for 2 m and stored at −70°C until further use.

### Cells

Vero cells (ATCC CCL-81, Manassas, VA) were used for titration and neutralization assays. Vero cells are African green monkey kidneys cells were maintained in DMEM with 5% FBS and 1% P/S and incubated at 37°C with 5% CO_2_.

### Infectious MAYV detections in inocula, RM, and mosquito samples

Infectious MAYV was quantified from RM inocula, plasma, and mosquito tissues using plaque assays on Vero cells. Mosquito bodies were initially tested for infection. If virus was detected, legs and wings were subsequently tested, following the route of virus dissemination in a mosquito [34,35]. If infectious virus was detected in legs and wings, saliva samples were passaged in Vero cells. Saliva was passaged to maximize virus detection as our prior CHIKV studies [32] show transmitted doses are low. For saliva, 50-75 µl of sample in DMEM was inoculated onto a single well of 60% confluent Vero cells in a 24-well plate. An additional 25-50 µl DMEM was added, and the plate was incubated for 1 h, with rocking every 10 min to facilitate viral attachment and maintain cell hydration. After incubation, 1 ml of DMEM was added to each well, and plates were incubated at 37°C with 5% CO_2_ for 3 days. If cytopathic effects (CPE) were observed 3 dpi in the test wells but not in the negative control wells, the supernatant was harvested and stored at −80°C. If no CPE was observed, the supernatant was transferred to a 1.5 ml tube, vortexed for 10 s, and then 100 µl was inoculated into fresh Vero cells for a second passage under identical conditions to the first passage. If CPE were detected in either passage, the saliva sample was considered positive for infectious MAYV. Inocula, RM tissues, plasma, and mosquito tissues were serially diluted and titrated. Vero cells were seeded into 12 (RM)- or 24 (mosquito)-well plates and incubated until 100% confluent. Plasma and homogenized tissue samples were diluted in ten-fold series in DMEM in technical duplicates, with dilution ranges of 10^-1^-10^-11^ (plasma), 10^-1^-10^-6^ (RM tissues), and 10^-1^-10^-12^ (mosquitoes). For infection assays, 125 µl (12-well) or 100 µl (24-well) of each diluted sample was inoculated per well and incubated for 1 h, with rocking every 10 m. After incubation, 2 ml (12-well) or 1 ml (24-well) 0.4% agarose (Genesee Scientific, El Cajon, CA) -DMEM at 42°C was added. Plates were incubated for 3 days, then fixed with 2% formalin for 1 h. The agarose overlay was removed and plates were stained with 0.05% weight/volume crystal violet (Millipore-Sigma, Burlington, MA) for 5 m, rinsed with deionized (DI) water, and dried before reading on a light box. Viral titers were calculated using the following equation:

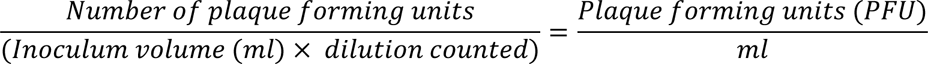

The infectious MAYV titer of a sample is presented as the mean of duplicate titrations. The limit of detection of titrations was 80 PFU/ml (12-well plate) or 10 PFU/ml (24-well).

### MAYV RNA quantitation by reverse transcription quantitative polymerase chain reaction (RT-qPCR)

MAYV RNA was extracted from all RM and mosquito samples using a viral RNA isolation kit and machine (MagMAX™, Waltham, MA). Two hundred microliters of sample were mixed with 10 µl RNA binding beads, 10 µl lysis binding enhancer, 120 µl 100% isopropanol (Geel, Antwerp, Belgium) and 120 µl lysis binding solution, which was loaded into the machine in a 96 deep well plate. Extracted RNA was eluted in 65 µl of Elution Buffer (ThermoFisher, Waltham, MA) for storage at −80°C prior to quantification. Into each well of the RT-qPCR plate, 10 µl of extracted RNA, 0.8 µl each of the 100mM forward and reverse primers, 0.2 µl of 100mM probe, 5 µl of TaqMan™ Fast Advanced Master Mix 4x (Applied Biosystems™, Waltham, MA), and 3.2 µl of nuclease-free water (Waltham, MA) were added. Each RT-qPCR plate included 2 wells of negative controls consisting of nuclease-free water (Waltham, MA), and 2 wells of each serial 10-fold positive control dilution that comprised standards of *in vitro* transcribed MAYV RNA derived from the infectious clone, ranging from 8.1 x 10^9^ to 8.1 x 10^-2^ genomes/ml, measured using a Qubit (Waltham, MA) prior to serial dilution. The RT-qPCR conditions on the Applied Biosystem™ ViiA 7 real-time PCR machine (Waltham, MA) were as follows: 52°C 15 s, 94°C 2 min, then 40 cycles of 94°C 15 s, 55°C 40 s, 68°C 20 s.

All samples tested by RT-qPCR used the following one of the follow primers and probe sets (all listed (5’→3’):

Set 1 (used for RM plasma and mosquito samples) Forward: AGCGATGAAAGGAGTACGCA (571→590) Reverse: TGGAACCTACCGCGAACATT (812→793) Probe: FAM-ACGAACAGGTGTTGAAAGCCAGGA-TAMRA (678 -> 701) previously published [28]

Set 2 (used for RM tissue samples) Forward: GGTAATGATCCACAGTCCATGC (5156→5177) Reverse: CGGGTTGAACAACCGGTTC (5237→5219) Probe: FAM-CTGACAACCAACAACCCATATCAACGG-NFQ-MGB (5190→5216)

Every sample was tested using technical triplicates. Both primer sets and the set 2 probe were designed *de novo* with primers and probes from ATCC (Manassas, VA). The limit of detection (LOD) of Set 1 was 8.1 x 10^4^ genomes/ml plasma or 8.1 x 10^5^ genomes/mosquito tissue, as determined by averaging the lowest RNA titer detected in the standards of all RT-qPCR assays. The LOD of Set 2 was 1.0 x 10^3.5^ genomes/gram or genomes/ml. Genomes/ml to genomes/mosquito tissue conversions were generated with the following calculations:

Mosquito tissues were saved in 500 µl DMEM / tissue, homogenized as described below, and 200 µl of the homogenate was place into the extraction reaction.

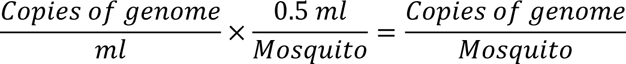

To convert PFU/ml to PFU/mosquito tissue, the PFU/ml titer was adjusted to account for the volume of media the mosquito tissue was homogenized in (500ul) and the fraction of that homogenate that was used for serial dilutions (12ul for 24-well plates, 2.4% of total volume) as so:

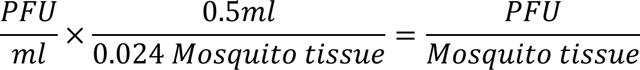

The MAYV RNA level in a sample is presented as the mean of triplicate measurements. Samples with no detectable RNA levels are presented at the LOD. The values for non-detectable samples are included in means as the LOD value.

### Neutralizing antibody quantification by plaque reduction neutralization test (PRNT)

MAYV neutralizing antibodies in RM serum were quantified using a PRNT_80_ assay in a 12-well format using MAYV strain IQT4235, the homologous strain used for RM inoculation. The PRNT_80_ titer was defined as the serum dilution that neutralized 80% of plaques compared to a virus-only control sample, which consisted of approximately 40 PFU of MAYV without RM serum. Sera were heat inactivated at 56°C for 30 min to destroy complement proteins. Controls included a positive serum from a MAYV-inoculated RM in this study that showed neutralization in a preliminary screening and a negative control consisting of serum from a RM with no alphavirus exposure history in a 1:100 final dilution. Each of these serum controls was tested with and without MAYV. A negative control of DMEM only was also included. All samples and controls were tested in technical duplicates. For the 12-well assays, serum was diluted 1:10 in technical duplicates, then serially two-fold diluted from 1:10-1:5120. Serum dilutions were mixed 1:1 with virus to make a final series of 1:20-1:10240. Serum-virus samples and controls were incubated at 37°C with 5% CO_2_ for 1 h. Following incubation, serum-virus mixtures or controls were inoculated onto plates of Vero cells at a volume of 125 µl/well and incubated for 1 h with rocking every 10 min. After incubation a 0.4% agarose-DMEM overlay at 42°C was then applied to wells. Plates were incubated for 3 days at 37°C with 5% CO_2_, then fixed with 2% formalin for 1 h. After removal of the agarose overlay, plates were stained with 0.05% weight/volume crystal violet for 5 min, washed with DI water, and dried before counting on a light box. PRNT_80_ titers are reported as the mean inverse serum dilution that resulted in 80% plaque reduction compared to virus-only control wells. If duplicate tests yielded different PRNT_80_ titers, the sample was re-tested, and all re-tested samples showed concordance in both replicates. The limit of detection for this assay was 20.

### MAYV deep sequencing from in vitro transcribed MAYV RNA, RM, and mosquito saliva samples

MAYV genomes were sequenced from *in vitro* transcribed RNA that was used to generate the virus stock that was inoculated into RM, plasma samples from all 12 RM at 2 dpi, and from all Vero passaged *Ae. aegypti* saliva samples that tested MAYV positive. Viral RNA from plasma was extracted a second time from the same samples previously tested by RT-qPCR following the same extraction protocol. Viral RNA from passaged saliva samples was extracted from Vero supernatants that had been thawed once. The extracted RNA was sequenced using ClickSeq [36]. To enrich and capture MAYV genomic RNA, we designed set of tiled MAYV primers (**Table 1**) that match with 100% identity to the MAYV genome used in these studies (GenBank: MK070491). 10 ul of extracted RNA was used as input in reverse transcription reactions, together with 10 pM of MAYV-specific tiled primers and mixture of dNTPs and 3′-azido-2′,3′-dideoxynucleotides (AzNTPs) (nucleotide analogues with modified 3′ groups that serve as chain terminators) at a ratio of 35:1, per the standard protocol for ClickSeq [37]. Primers were annealed by heating the extracted RNA to 95°C for 2 minutes, followed by slow cooling to 50°C. Reverse transcription components (Superscript IV, ThermoFisher, Waltham, MA) were added at 50°C to initiate cDNA synthesis. The reverse transcription reaction was incubation for 10 minutes at 50°C and then 10 minutes at 80°C, per the manufacturer’s standard protocol. 2.5 units of RNaseH were then added and incubated at 37°C for 20 minutes and then heat-denatured at 80°C for 10 minutes. The 3′-azido-blocked cDNA fragments were then “click-ligated” to the 5′-hexynyl adaptor containing the full Illumina i5 adaptor with i5 indexes via copper(I)-catalyzed azide-alkyne cycloaddition (CuAAC). Click-ligated cDNA was used as input to a PCR reaction that provided i7 indexes for 21 cycles. cDNA library fragment sizes of 300– 600 nt were selected using double-sided SPRI bead purification and sequenced on an Illumina NovaSeq X, yielding >10M 150x2 paired end reads per sample. Raw FASTQ data were preprocessed (adaptor trimming and quality filtering) using *fastp* [38] and *cutadapt* [39], using batch scripts previously published for analysis Tiled-ClickSeq data [40]. Reads were then mapped to the MAYV reference genome (GenBank: MK070491) with *bowtie2* [41] and single nucleotide variants and minority variants were called using *pilon* [42].

**Table 1.**
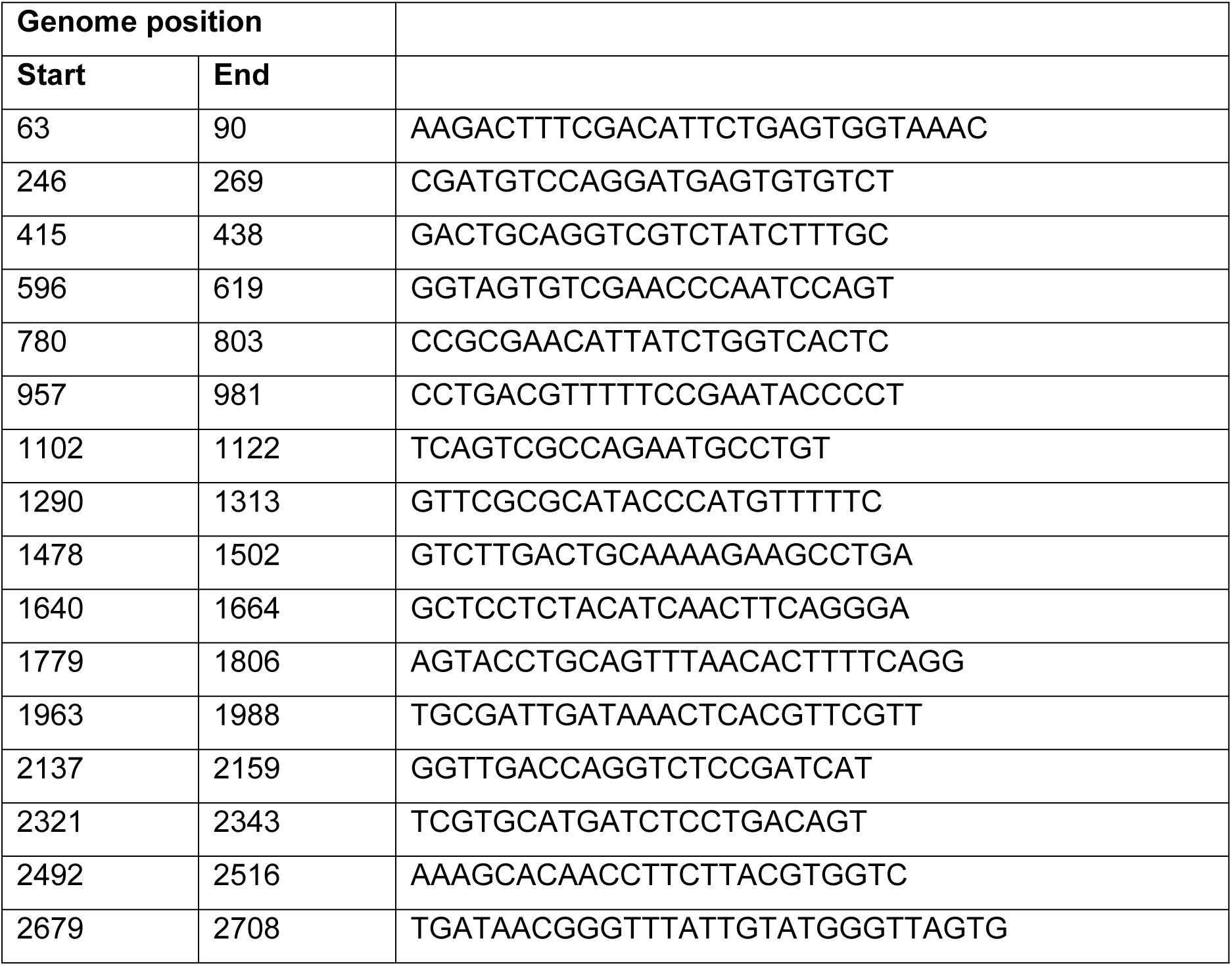
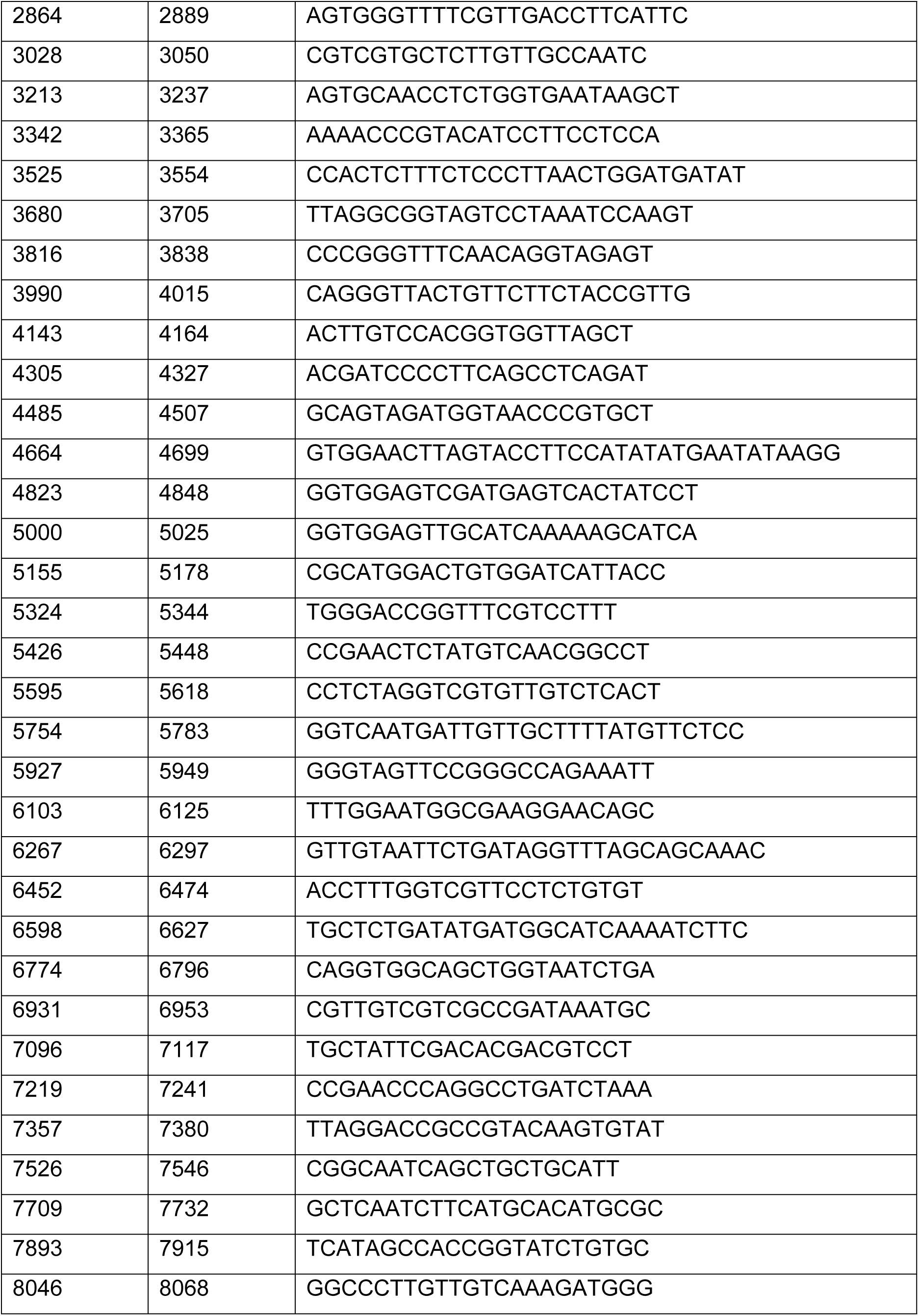
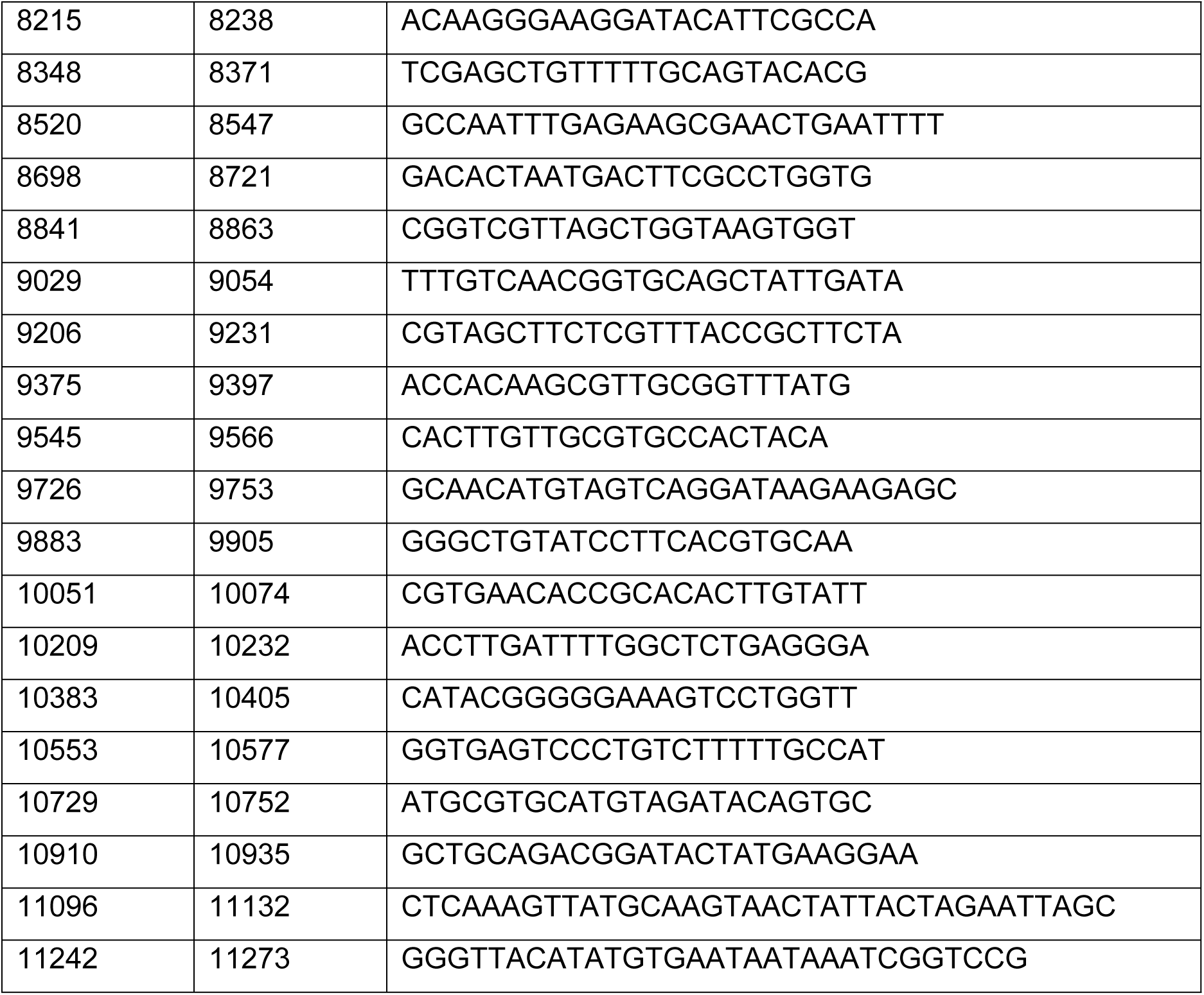
Tiled MAYV primers used for library generation for sequencing. Primers were designed to match the genome of the parent strain, GenBank: MK070491.

### Statistical Analyses

GraphPad Prism 10.3.1 was used for all statistical analyses, which are enumerated in the results. Analyses of infectious MAYV titers (PFU) and MAYV RNA were performed on log-transformed data.

### Data availability

Raw data are available as **Supplemental File 1**.

## RESULTS

### Rhesus macaques develop a 3-day MAYV infectious viremia

This study aimed to characterize infection dynamics and disease outcomes following inoculation of MAYV IQT4325 from Iquitos, Peru in RM, using 2 doses and routes of administration (**Figure 1A**). To evaluate the impact of inoculation route and dose on viremia kinetics, RM were randomly assigned to 3 cohorts of 4 animals each: 7 log_10_ PFU IV, 7 log_10_ PFU SC, and 3 log_10_ PFU SC. The IV route was chosen to ensure infection, as these studies were initiated before publication of recent MAYV NHP studies [11,12], when infectivity of MAYV IQT4235 for RM was not certain. The SC route was selected to mimic mosquito-borne transmission [43]. The 7 log_10_ dose exceeds mosquito-delivered doses and was intended to ensure infection. The 3 log_10_ dose simulated mosquito-delivered doses, where our prior studies with CHIKV [32] and studies by us [27] and others [28,29] for MAYV show that *Ae. aegypti* salivate between 1.0-2.5 log_10_ PFU or 1.0-4.0 log_10_ genomic RNAs into capillary tubes. Back-titration of the inocula confirmed that the administered doses were within 10% of the target doses. Clinical signs of disease were monitored, and blood samples were collected daily from 1-7 dpi and at necropsy, 10 or 12 dpi. Infectious MAYV and MAYV RNA were quantified from plasma, while tissues were harvested and analyzed for MAYV RNA levels. Histopathological evaluation of muscle and joint tissues was conducted, with inflammation scored using a quantitative metric.

**Figure 1.**
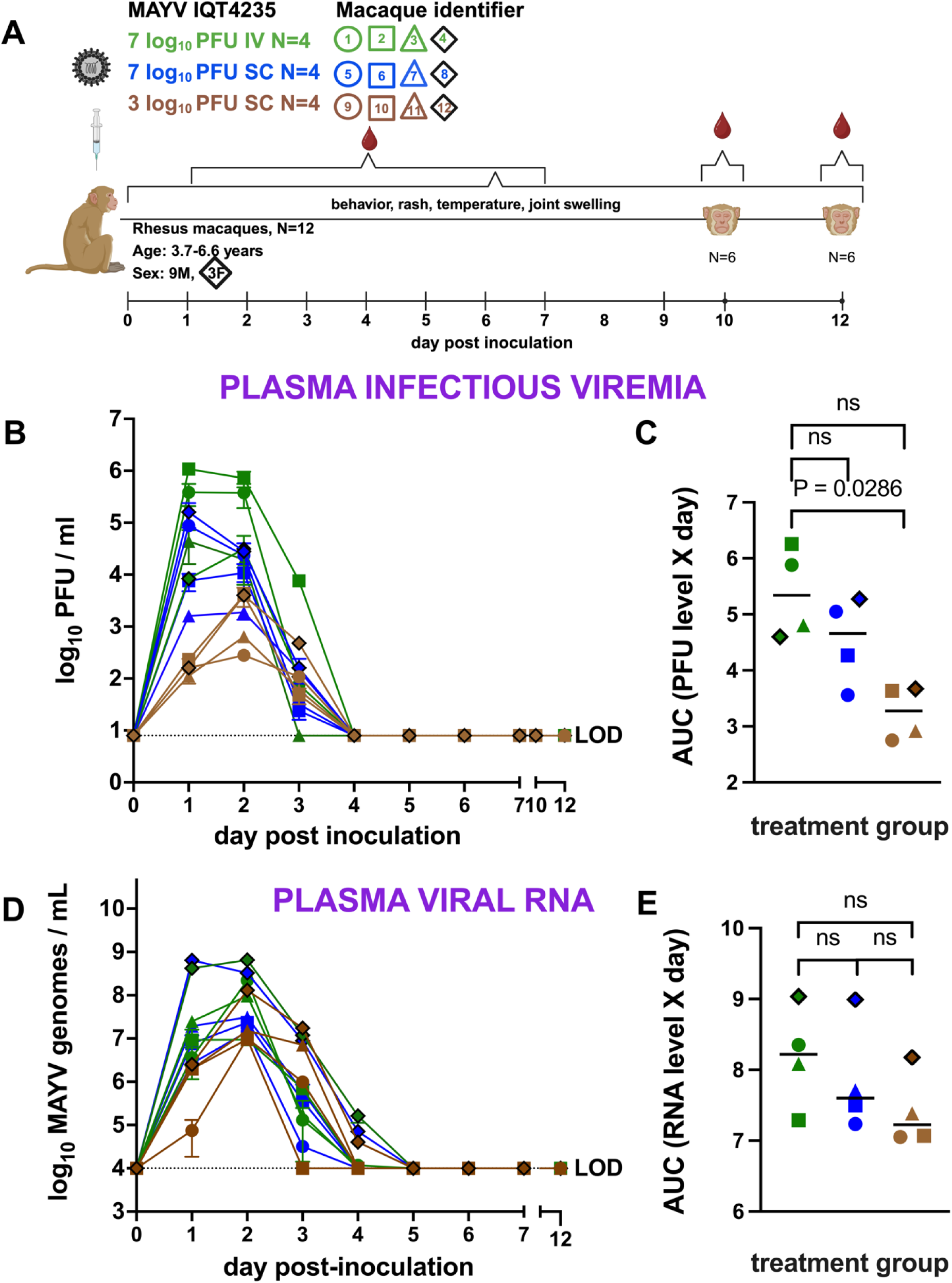
Rhesus macaques develop short MAYV viremias. (**A**) **Rhesus macaque study design** in which 12 rhesus macaques (9 males and 3 females) were inoculated with 3 or 7 log_10_ PFU MAYV by intravenous (IV) or subcutaneous (SC) routes on day 0, followed by blood collections 1-7 days post-inoculation (dpi) and euthanasia 10 or 12 dpi. (**B**) **MAYV infectious viremia** in plasma 1-12 dpi measured as plaque forming units (PFU)/ml. Each line shows kinetics from 1 RM and bars report standard deviations from triplicate titrations. (**C**) **Area under the viremia curve** (AUC). Horizontal lines denote averages. **D**) **MAYV RNA in plasma** measured as genomes/ml 1-7 and 10 or 12 dpi. Each line shows kinetics from 1 RM and bars report standard deviations from triplicate RT-qPCR measurements. (**E**) **Area under the MAYV RNA curve**. Horizontal lines denote averages. Data points outlined in black represent female animals. P values are based on two-tailed Mann-Whitney tests, ns = not significant (p>0.05). The colors and shape identifiers denote individual RM, which are also identified with a unique number. LOD is the limit of detection.

None of the MAYV-inoculated RM exhibited signs of behavioral changes or joint swelling, but 8 animals exhibited other signs of mild clinical disease between 1 and 4 dpi. Three animals had mildly elevated temperatures, 3 animals had nausea, 5 animals developed mild erythema at the inoculation site, and one animal developed an abdominal rash prior to mosquito feeding. Some of the rashes were detected on a day of mosquito feeding and on the shaved area of the abdomen where mosquitoes were presented. Eight of 12 RM also developed rashes or skin redness in a region distant from the abdomen and not on a day of mosquito presentation (**Supplemental Table 2**). All MAYV-inoculated RM developed detectable viremia, with infectious MAYV in plasma observed from 1-2 (N=1) or 1-3 (N=11) dpi, peaking at 2-6 log_10_ PFU/ml. No infectious MAYV was detectable in plasma after 3 dpi (**Figure 1B**). Peak infectious viremia occurred at 1 (N=4) or 2 (N=8) dpi, and mean infectious viremia between the 1 female and 3 male RM in each group were similar, although statistical analyses were not performed given only 1 female was in each group. RM that received 7 log_10_ PFU IV had significantly higher mean viremia areas under the curve (AUC) compared to those receiving 3 log_10_ PFU SC (Mann-Whitney, p=0.02) (**Figure 1C**). However, no significant difference in AUC was observed between the 2 SC groups. The detection pattern of MAYV RNA in plasma mirrored that of infectious virus, with RNA detected from 1-3 (N=8) and 1-4 dpi (N=4) animals, peaking at 7-9 log_10_ genomes/ml (**Figure 1D**). MAYV RNA levels peaked 1 (N=1) or 2 (N=11) dpi, and in most samples, MAYV RNA levels were typically several logs higher than infectious MAYV levels.

There were no significant differences in mean genome AUC levels across cohorts (Kruskal-Wallis, p>0.05) (**Figure 1E**). Together, these results demonstrate that RM are susceptible to MAYV strain IQT4235 with mild clinical disease and focal rash in some animals. Viremia was short-lived, with infectious virus becoming undetectable after 3 dpi and viral RNA levels undetectable after 4 dpi. No sex differences in viremia kinetics were observed. Data from different dose and route groups show that 7 log_10_ PFU IV results in higher infectious viremia AUC compared to 3 log_10_ PFU SC but viremia and viral RNA kinetics were similar in RM receiving 3 versus 7 log_10_ MAYV via SC injection.

### MAYV-infected rhesus macaques develop neutralizing antibody responses

To investigate the humoral immune response following experimental MAYV inoculation, we measured endpoint neutralizing antibody titers in RM serum upon euthanasia, 10 or 12 dpi, against the homologous strain of MAYV used for RM inoculation. All 12 animals developed detectable PRNT_80_ titers, ranging from 80 to 1280 (**Figure 2A**), with a mean titer of 160. There was no significant difference in mean titers between the different dose and route groups (Kruskal-Wallis, p>0.05).These data demonstrate RM generate MAYV-specific neutralizing antibody responses within 10 to 12 days of MAYV infection.

**Figure 2.**
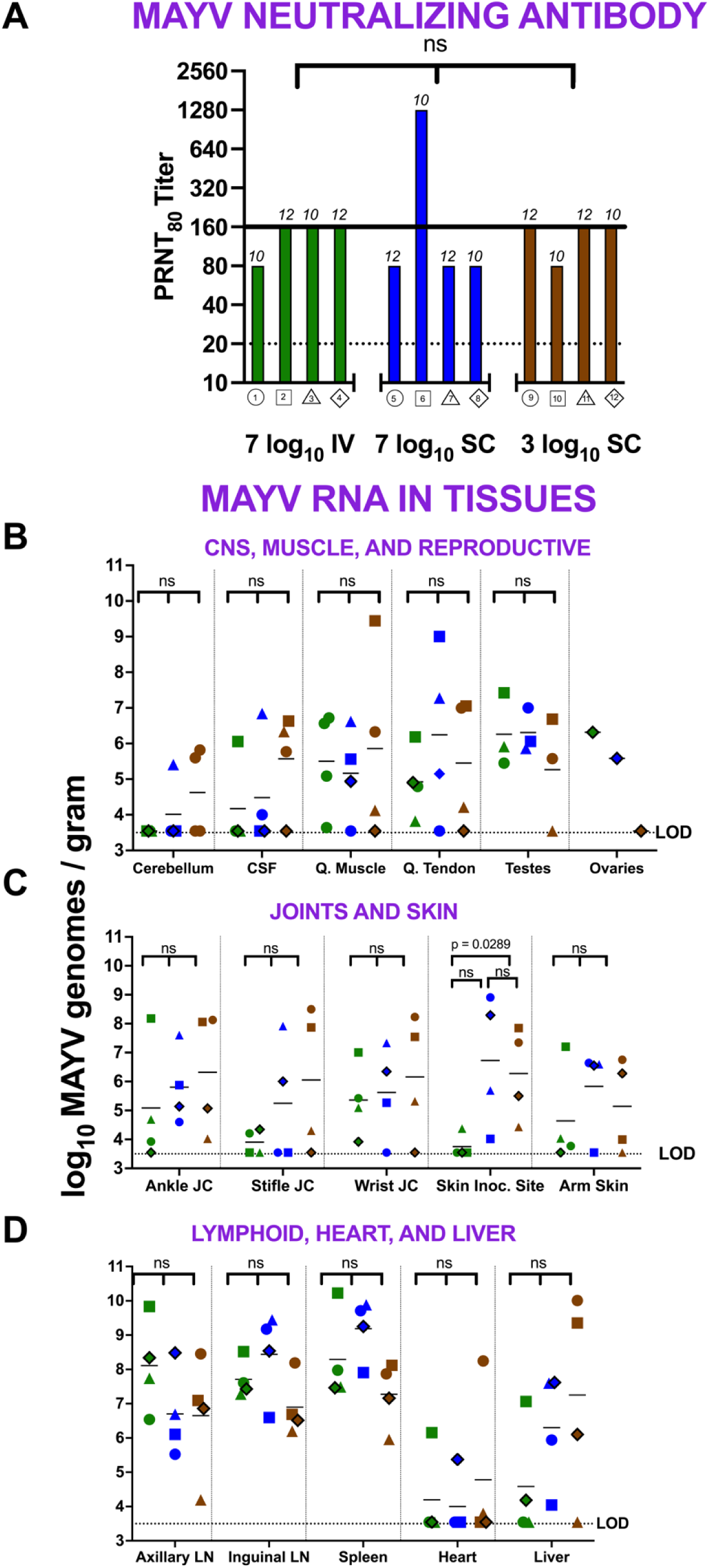
Rhesus macaques develop MAYV neutralizing antibodies in serum and MAYV RNA in multiple tissues 10 or 12 dpi. (**A**) **Plaque reduction neutralization test 80%** (**PRNT**_80_**) titers in serum.** PRNT_80_ titers are based on duplicate titrations. Numbers above bars show day of euthanasia on which PRNT_80_ titers were measured. The solid black line shows the average titer for all RM. The colors and shape identifiers denote individual RM, which are also identified with a unique number. The dotted line shows the LOD. **MAYV RNA levels in** (**B**) **central nervous system (CNS), quadriceps, and reproductive organs** and (**C**) **joints and skin,** and (**D**) **lymphoid tissues, heart, and liver**. The colors and shapes identify individual RM. Data points with black outlines represent female animals. MAYV RNA levels were measured by RT-qPCR with an average LOD of 3.5 log_10_ genomes/gram, indicated by the horizontal dotted line. Cerebrospinal fluid data is graphed in genomes/ml with the same LOD. Horizontal lines show mean titers. Values from animals without detectable MAYV RNA are reported at the LOD and are included in means. “ns” indicates no statistically significant difference by Kruskal-Wallis test, p>0.05. The p-value shown above “Skin Inoc. Site” is based on a Mann-Whitney test. “CSF” = cerebrospinal fluid; “Q. Muscle/Tendon” = quadricep; “JC” = joint capsule; “inoc.” = inoculation; “LN” = lymph node.

### MAYV RNA tropism is diffuse and includes muscle, lymphoid, central nervous, and circulatory systems of rhesus macaques

We evaluated whether infectious MAYV or MAYV RNA could be detected in various tissues, including muscle and joint, as well as other systems previously identified as targets in male rhesus [12] and female cynomolgus [11] macaques. We measured both infectious MAYV and MAYV RNA levels in 14 non-reproductive tissues for each RM and in testes and ovaries. No infectious MAYV was detected in any of the RM tissues.

However, MAYV RNA was detected in every tissue type for at least one RM (**Figure 2B-D**). We observed considerable variability in MAYV RNA detection both within and across cohorts and tissue types. The skin inoculation site in the 3 log_10_ PFU SC group had significantly higher mean MAYV RNA titers compared to the 7 log_10_ PFU IV inoculated group (Mann-Whitney, p=0.02). No other statistically significant differences in mean tissue titers were observed across groups (Kruskal-Wallis, p>0.05).

The mean MAYV RNA level across all tissues ranged from 3.5-10.0 log_10_ genomes/gram. Most RM in each group had detectable MAYV RNA in the majority of their tissues (**Figure 3**). For the reproductive tissues, all but one male and one female had detectable MAYV RNA in their testes and ovaries, respectively. These data show that MAYV infects multiple RM tissue systems, as evidenced by viral RNA detections in most animals at 10 or 12 dpi, nearly a week after MAYV RNA clearance from the blood. In general, the highest MAYV RNA levels were detected in lymphoid tissues including lymph nodes and spleen.

**Figure 3:**
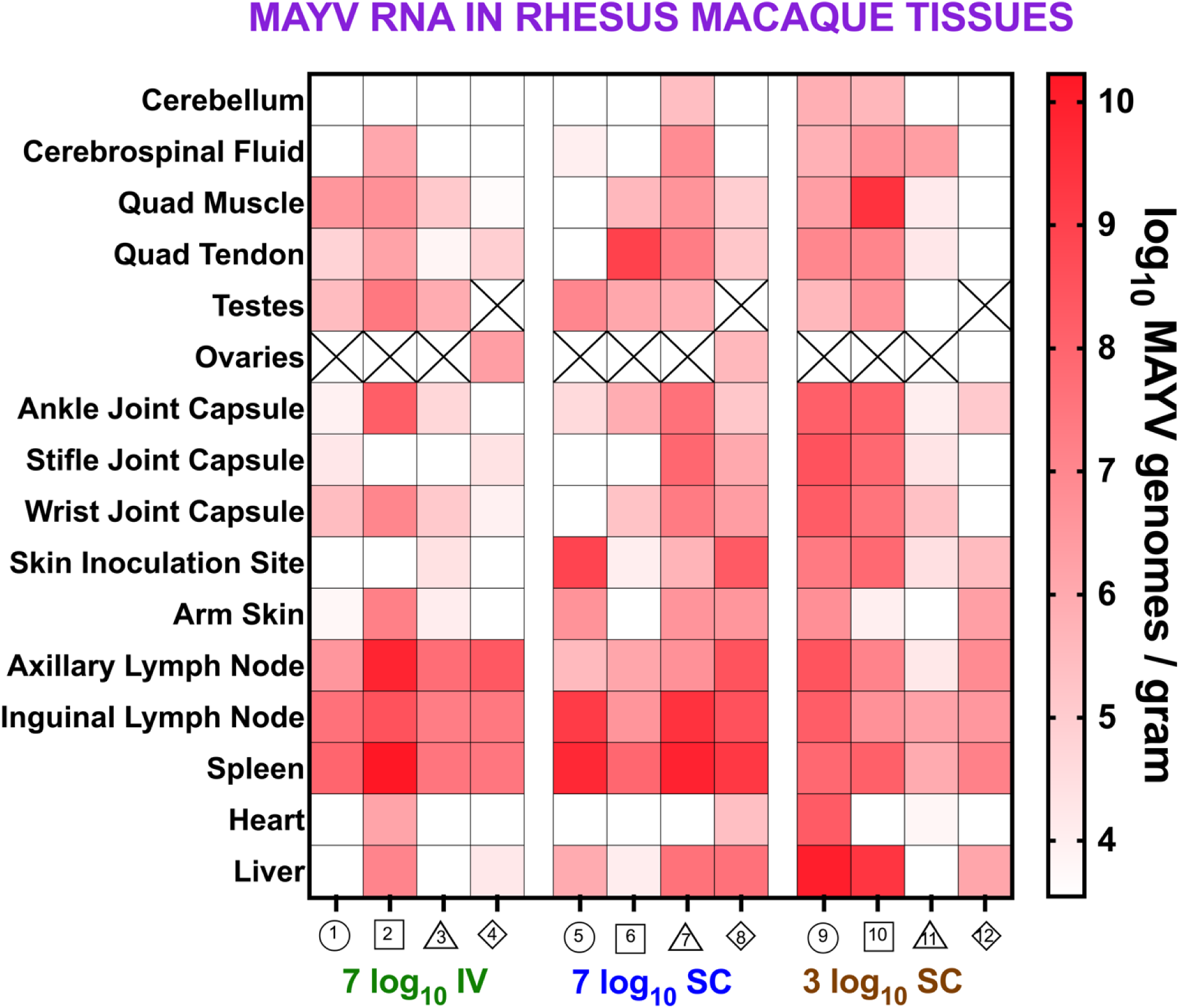
MAYV RNA is detectable in multiple tissues of rhesus macaques 10 or 12 dpi. The heatmap shows the magnitude of MAYV RNA detection. The colors and shape identifiers denote individual RM, which are also identified with a unique number. Female animals (4, 8, and 12) are represented as diamonds. MAYV RNA levels were measured by RT-qPCR with an average limit of detection (LOD) of 3.5 log_10_ genomes/gram. Boxes with X symbols show tissues that were not available.

### MAYV infection of rhesus macaques produces histopathologic signs of inflammation in in joints and muscles

After establishing the susceptibility, viremia kinetics, and tissue tropism of MAYV IQT4235 in RM, we focused on identifying histopathologic changes in target tissues. Since alphaviruses like MAYV target joints and muscles, causing muscle pain and arthritic disease, we concentrated on these tissue types. Using our quantitative histopathologic scoring scale (**Supplemental Table 1**), we identified and quantified inflammatory changes in hematoxylin and eosin-stained joint and muscle tissues. Inflammation was considered present if any of the 12 tissues evaluated showed had a score greater than 0, where 0 represents the baseline for colony control animals not exposed to MAYV. All 12 MAYV inoculated RM showed histopathologic signs of inflammation, with joint scores ranging from 6 to 14 and muscle scores from 1 to 10 (**Figure 4A**). No significant differences were found in mean pathology scores in joint, muscle, or composite (joint + muscle) tissues across any dose or route treatment groups (Kruskal-Wallis, p>0.05). Additionally, there was no significant difference in mean pathology scores between RM euthanized at 10 or 12 dpi (t-test, p>0.05). Representative images of the inflammatory responses show immune cells (dark purple, marked with asterisks*) in the joint synovium for a finger (**Figure 4B**), toe (**Figure 4C**), and elbow (**Figure 4D**). These data demonstrate that MAYV caused histopathologic signs of muscle and joint inflammation in all infected RM at 10 or 12 dpi. Furthermore, the route of inoculation and dose did not affect the magnitude of these histopathologic changes at these sites.

**Figure 4.**
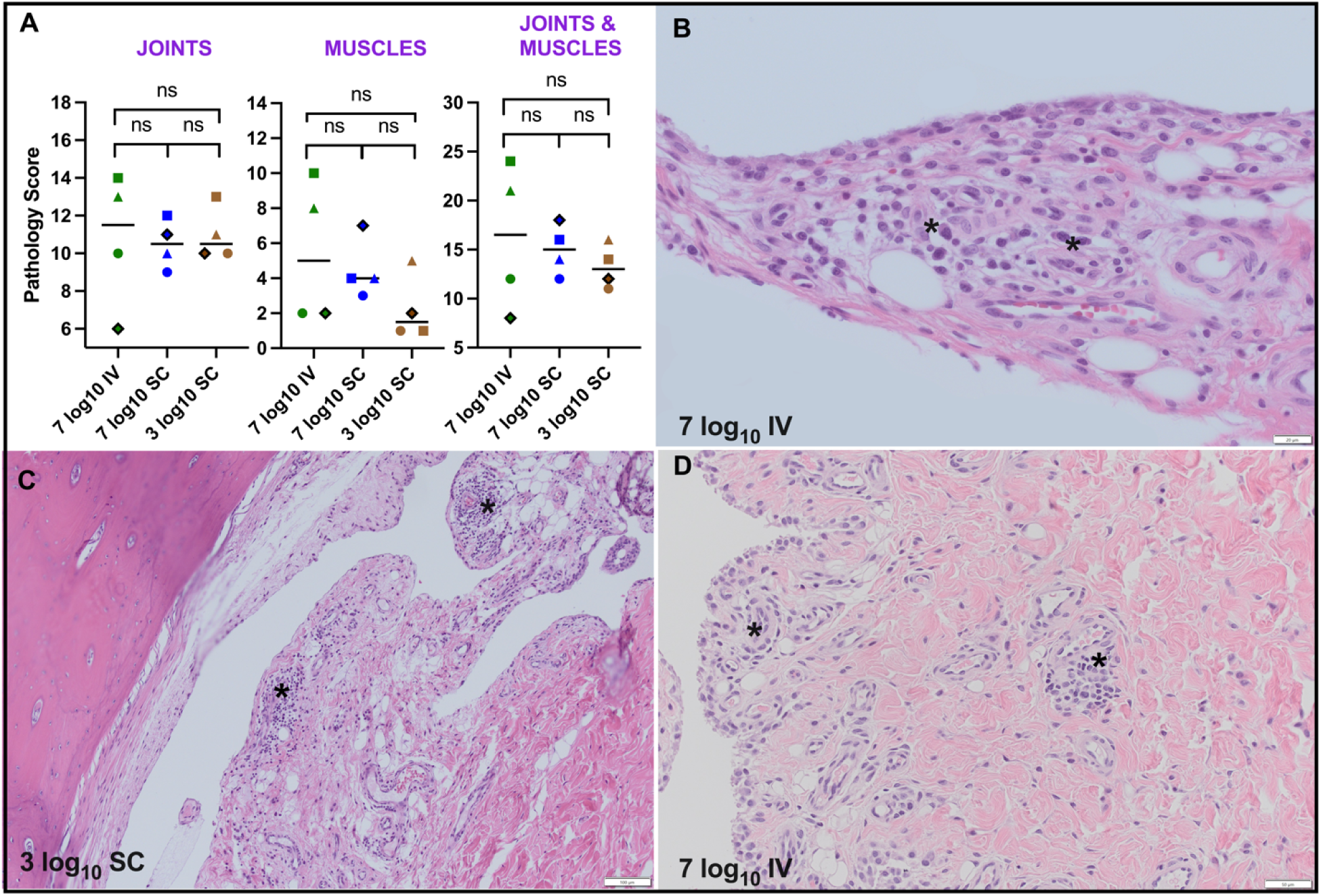
MAYV infection of rhesus macaques produces mild inflammatory responses in joints and muscles. (**A**) **Pathology scores** in joints and muscles for MAYV infected RM based on quantitative histopathologic criteria (**Supplemental Table 1**). The colors and shape identifiers denote individual RM. Bars show means. ns denotes not significant (Kruskal-Wallis, p>0.05). Representative histology images of (**B**) **Joint synovium of finger**, animal #2 from 7 log_10_ PFU IV group, scale bar: 20 micrometers. **(C) Joint synovium of toe**, animal #10 from 3 log_10_ PFU SC group, scale bar: 100 micrometers. **(D) Joint synovium of elbow,** animal #2 from 7 log_10_ PFU IV group scale bar: 50 micrometers. Asterisks indicate immune cell(s).

### MAYV-viremic rhesus macaques infect *Ae. aegypti* 2 days after macaque inoculation

To evaluate the ability of RM to transmit MAYV to a primary arbovirus vector in urban settings, we allowed female *Ae. aegypti* mosquitoes, sourced from a colony originating from Los Angeles, California, USA, to bloodfeed on each anesthetized RM 2, 3, 5, and 7 dpi (**Figure 5A**). These times were chosen based on our projection of when the animals would be viremic. Since we did not test RM plasma in real-time, we were unaware that animals cleared detectable viremia after 3 dpi, and neither of the 2023 macaque studies had been published at the time our study was conducted. All mosquitoes that bloodfed were incubated for 10 days, the period during which MAYV disseminates and reaches mosquito saliva [27–30]. We assessed MAYV infection in mosquitoes by testing their bodies, legs and wings for dissemination, and saliva for transmission. All RM showed detectable viremias at 2 and 3 dpi, but none were viremic at 5 or 7 dpi (**Figure 5B**). The timing of mosquito presentations therefore covered both viremic (2 and 3 dpi) and aviremic (5 and 7 dpi) periods in RM. Except for one mosquito that fed 3 dpi, only RM 2 dpi were infectious to *Ae. aegypti* (**Figure 5C**). For mosquitoes that fed 2 dpi, 48% (104/217) of bodies tested positive for MAYV. Among these, 38% (39/104) of their legs and wings tested positive. These data demonstrate that MAYV-infected RM can infect *Ae. aegypti* but the window of infectiousness is narrow and does not extend past 2 dpi. Dissemination rates from these mosquitoes were also low.

**Figure 5.**
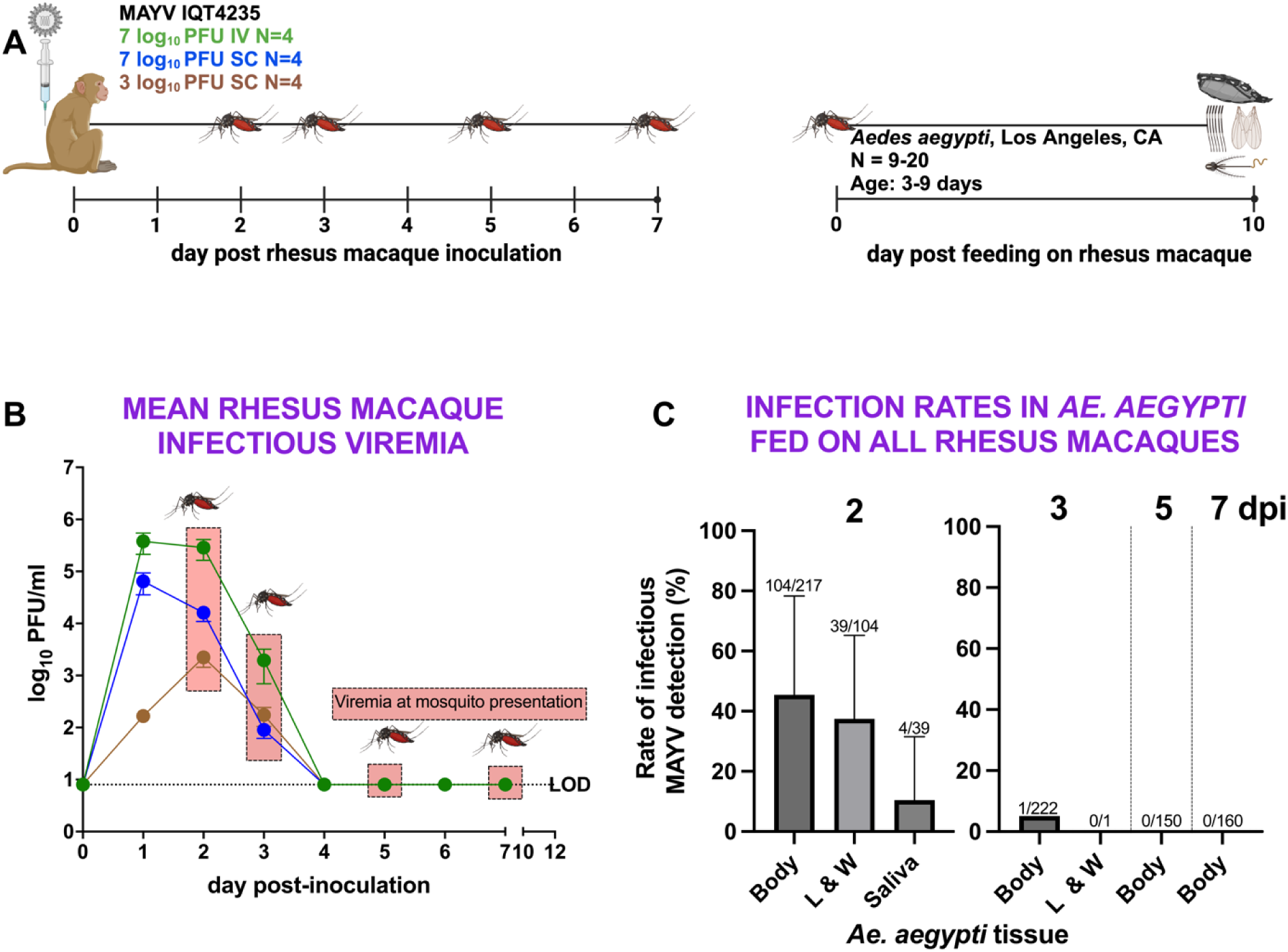
MAYV viremic rhesus macaques are infectious to *Ae. aegypti* mosquitoes in a narrow window, 2 dpi. **(A) Experimental design** showing the timing of mosquito exposure to MAYV-infected RM 2, 3, 5, and 7 dpi, followed by mosquito incubation and harvest of bodies, legs and wings, and saliva 10 days post-feed (dpf). (**B) Mean viremia levels in RM on mosquito feeding days**, represented as cohort log-transformed means from the data shown in Figure 1B. Error bars show standard deviations. **(C) Rates of MAYV infection in *Ae. aegypti*** (bodies), dissemination in legs and wings (L&W), and transmission in saliva from mosquitoes that fed on RM 2, 3, 5, or 7 dpi. Error bars show standard deviations and the fractions above each error bar represent the number of mosquitoes with detectable infectious MAYV divided by the total number of mosquitoes tested.

### Few *Ae. aegypti* that feed on viremic rhesus macaques transmit infectious MAYV

To evaluate the transmissibility of MAYV by infected mosquitoes, we used detection of infectious virus in saliva as a surrogate for transmission to a vertebrate. Saliva samples from individual *Ae. aegypti* were tested using a sensitive Vero cell passaging approach. Saliva collected into capillary tubes from mosquitoes with detectable infectious MAYV in their legs and wings was passaged once on Vero cells. Samples showing cytopathic effects in Vero cells were considered to contain infectious MAYV. Of the 39 mosquitoes with disseminated infections in legs and wings, 4 (10%) had infectious MAYV in saliva. Of these four mosquitoes, one fed on RM 3, two fed on RM 4 and one fed on RM 12. The overall potential transmission rate for all 217 *Ae. aegypti* that fed on RM in this study was 2% (4/217), and 4% (4/104) of mosquitoes with infected bodies. These data show that while *Ae. aegypti* infected from RM can transmit MAYV, such transmission is rare.

### Individual rhesus macaques are variably infectious to *Ae. aegypti*

To assess MAYV infectivity in *Ae. aegypti* that ingested MAYV from viremic RM 2 dpi, we quantified infectious MAYV in bodies (representing infection) and legs & wings (representing disseminated infection) 10 dpf. The mean infection rates were significantly higher in cohorts that fed on RM inoculated with 7 log_10_ PFU compared to those that fed on the 3 log_10_ PFU cohort (unpaired t-test, p<0.001). However, there was no significant difference in mean infection rates between the 2 high-dose groups or the 2 SC groups (unpaired t-test, p>0.05). (**Figure 6A**). Eleven of 12 RM infected at least one *Ae. aegypti* body. Infection and dissemination rates were varied widely within cohorts that fed on different RM, ranging from 5% (1/20) to 90% (18/20) for infection rates, and 0% (0/1) to 100% (4/4) for dissemination rates (**Figure 6B**).

**Figure 6:**
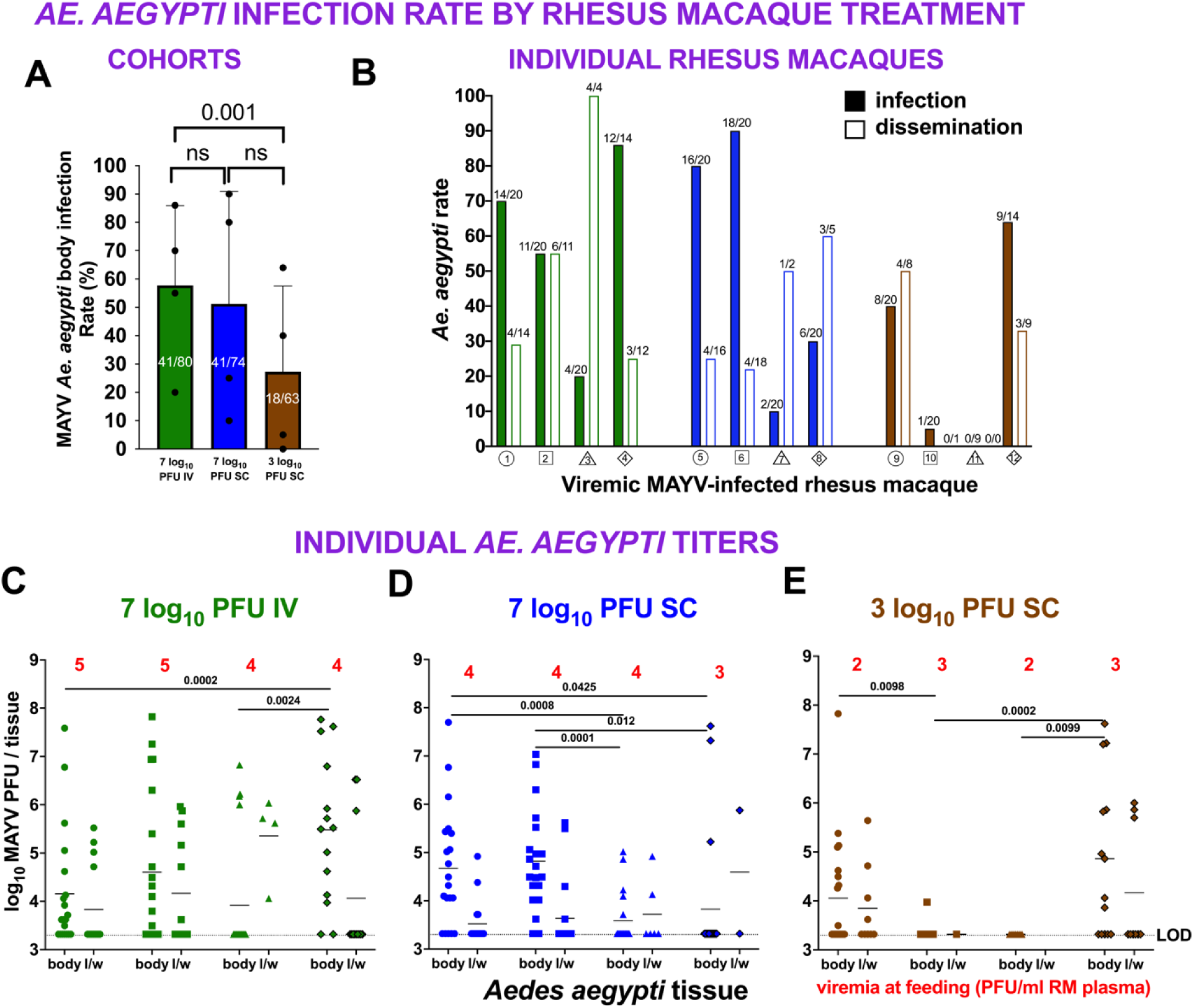
MAYV infects and disseminates *Ae. aegypti* that ingest bloodmeals from viremic rhesus macaques 2 dpi. **(A) Cohort infection rates** of *Ae. aegypti* exposed to RM treated with the same dose and route of MAYV. Each dot represents the mean infection rate from a cohort that fed on a single RM, with error bars indicating standard deviations. Fractions on bars represent number of positive mosquito bodies divided by the number of bloodfed mosquitoes. **(B) Rates of infection** (bodies, solid bars) and **dissemination** (legs and wings, open bars) in cohorts of *Ae. aegypti* that ingested bloodmeals from each viremic RM. Fractions above bars show the number of positive tissue samples divided by the number tested. The colors and shape identifiers denote individual RM, which are also identified with a unique number. **(C-E) Infectious MAYV levels in bodies and legs of individual *Ae. aegypti*** that ingested bloodmeals from viremic RM. Doses and routes of RM treatment are indicated by colored data points, and red numbers show the plasma MAYV titers at the time of mosquito feeding (2 dpi). Each dot represents a single mosquito tissue titration, with lines showing the mean values. The horizontal dotted line shows the assay limit of detection (LOD), 3.3 log_10_ PFU/tissue. The colors and shapes of data points follow designations for individual RM, and data points with black outlines represent mosquitoes that fed on female RM.

### *Ae. aegypti* that feed on viremic rhesus macaques develop a range of MAYV levels

Mean *Ae. aegypti* body titers were significantly higher in the 7 log_10_ PFU IV cohort compared to the 3 log_10_ PFU SC cohort (unpaired t-test, p=0.01) but were not different between the two 7 log_10_ or two SC groups (unpaired t-test, p>0.05) (**Figure 6C-E**). Some mosquito cohorts from all three groups showed significantly higher mean titers than others within the same dose and route group (unpaired t-tests, p≤0.05). *Ae. aegypti* developed a range of MAYV titers, with cohort means ranging from just above the limit of detection of 3.3 to 5.5 log_10_ PFU/body and almost 8.0 log_10_ PFU/body in some individuals. For legs and wings, mean titers ranged from just above 3.5 to 5.4 log_10_ PFU/legs and wings and more than 6.0 log_10_ PFU/legs and wings in some individuals. The results from these studies demonstrate individual RM are variably infectious to *Ae. aegypti* 2 dpi, with the majority of animals infecting mosquitoes. Individual mosquitoes that fed on the same RM showed variable rates of infection and dissemination and variable magnitudes of infection.

### Rhesus macaque MAYV viremia at the time of mosquito feeding does not predict *Ae. aegypti* infection rates

The variable infectivity of RM for *Ae. aegypti* was accompanied by a 3 log_10_ viremia range (2-5 log_10_ PFU/ml blood, as indicated by the red values in **Figure 5C-E**) at mosquito presentation. This prompted us to investigate whether RM viremia at the time of mosquito feeding correlates with mosquito infection rates. We predicted that, similar to conventional vector competence studies that use virus-spiked bloodmeals, *Ae. aegypti* infected by viremic RM would show a dose-dependent response to MAYV infection. To test this, we performed a linear regression analysis comparing infection rates in *Ae. aegypti* that fed on MAYV-infected RM with those that fed on artificial virus-spiked bloodmeals (**Figure 7A**). The bloodmeal data came from a published study that used the same strain of MAYV and *Ae. aegypti* sourced from Iquitos, Peru [27]. The correlation between RM viremia and *Ae. aegypti* infection rate was low, with a coefficient of determination (R^2^) of 0.18 (**Figure 7B**). In contrast, the artificial bloodmeal data showed a much stronger correlation, with a R^2^ of 0.72 (**Figure 7C**). These results indicate that higher MAYV RM viremia does not predict increased *Ae. aegypti* infectiousness, contrasting with the dose-response patterns observed in artificial bloodmeal experiments.

**Figure 7.**
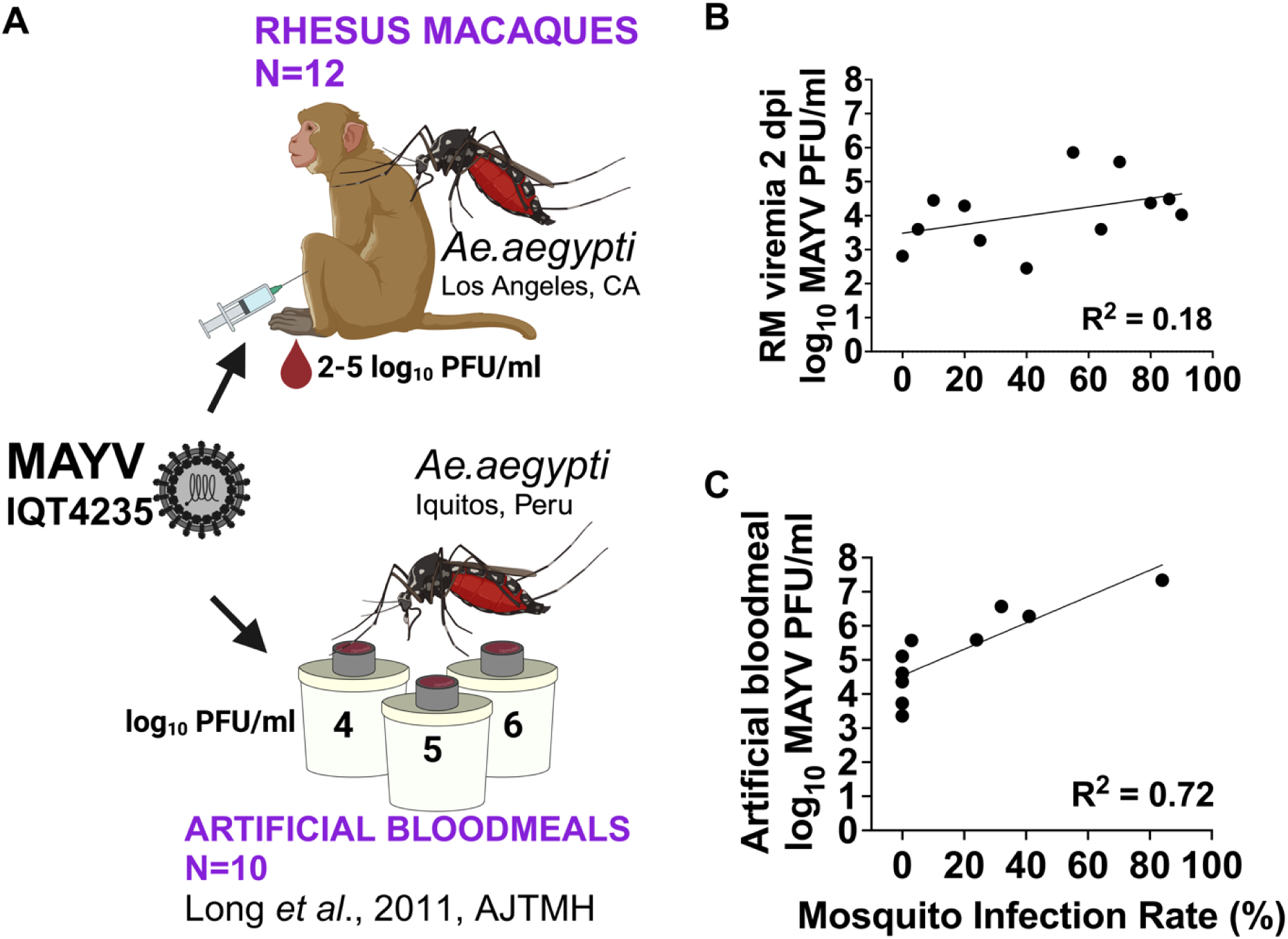
Rhesus macaque MAYV viremia levels do not correlate with *Ae. aegypti* infection rates, contrasting with the dose-response observed in studies using artificial bloodmeals spiked with virus. **(A) Comparison of MAYV infection sources**: *Ae. aegypti* fed on either RM (this study) or artificial bloodmeals based on prior published data [27]. Both studies used the same strain of MAYV, with *Ae. aegypti* sourced from Los Angeles, CA (this study) or Iquitos, Peru (published study). **Linear regression analysis** comparing **(B)** RM viremias at 2 dpi and infection rates in *Ae. aegypti* or (**C**) artificial bloodmeal MAYV titers and *Ae. aegypti* infection rates in cohorts that fed on bloodmeals of varying titers. The R^2^ value represents the coefficient of determination.

### Mayaro virus is genetically stable during rhesus macaque infection and transmission from *Ae. aegypti*

To evaluate whether genetic changes in MAYV arising during infection of RM or *Ae. aegypti* may influence infectivity of RM to mosquitoes which could explain the lack of correlation between the magnitude of RM viremia and *Ae. aegypti* infection rates, we deep sequenced the MAYV genomes in plasma from all 12 RM at 2 dpi and compared it to the MAYV plasmid sequence that was used to generate the virus stock that was inoculated into RM. We also sequenced all 4 Vero passaged MAYV positive *Ae. aegypti* saliva samples. All 12 plasmas and 3 of the 4 *Ae. aegypti* saliva samples were identical across the genome at the consensus level (>50% of RNAs) to the sequence of MAYV used to produce the virus stock for RM inoculations. There was a single non-coding mutation (G5758A) in 1 Vero passaged mosquito saliva sample from a mosquito that fed on RM 3. These data do not support viral genetic differences as a contributor to the variable infectiousness of RM for *Ae. aegypti*.

## DISCUSSION

This is the fourth study to investigate experimental MAYV infection of macaques, following the three previous studies that demonstrated susceptibility, tropism, and mild disease [10–12]. In this study, we inoculated 12 RM with 2 doses of MAYV and observed infectious viremias for 2-3 days, peaking at 2-6 log_10_ PFU/ml, along with 3-4 days of detectable plasma viral RNA. The higher area under the viremia curve in the 7 log_10_ PFU IV group compared to the 3 log_10_ PFU SC group suggests that both the inoculation dose and route influence the magnitude and kinetics of MAYV viremia in RM. All RM developed MAYV specific neutralizing antibodies, viral RNA in multiple tissues, and mild inflammation in the evaluated joints and muscles. With the limitation that group sizes were small, there were no significant differences in neutralizing antibody titers, joint and muscle histopathology scores, or tissue tropism among groups treated with different MAYV doses or inoculation routes. We observed similar MAYV viremia kinetics, tissue tropism, antibody levels, and joint and muscle inflammation in female and male RM, suggesting that sex does not affect MAYV infection or disease.

While existing murine models of MAYV disease replicate several features of Mayaro fever in people, such as weight loss, arthralgia, and elevated pro-inflammatory cytokines like monocyte chemoattractant protein (MCP)-1 and interferon (IFN)-gamma (γ) [4,44–46], NHP more accurately mimic humans due to closer genetic, physiological, anatomical, and immune system similarities. NHP models are also crucial for the licensure of viral vaccines [47]. NHP are particularly valuable for understanding human MAYV, as widespread outbreaks have not been well-documented, limiting understanding of clinical Mayaro fever. In humans, MAYV viremias at the time of clinic presentation range from 2.5-5.0 log_10_ PFU/ml [2,27] and are accompanied by fever, rash, and chronic arthralgia in up to 50% of cases [1,48,49]. The first published NHP study of MAYV, conducted in 1967, involved 6 RM inoculated SC with 4.7-5.9 log_10_ suckling mouse intracerebral inoculation lethal dose 50%. These animals developed 4-5 day viremias peaking at 3.0-5.3 log_10_ PFU/ml [10]. In 2023, Weber *et al*. [12] inoculated 3 male RM SC with 5 log_10_ PFU/ml MAYV BeAr 505411, a lineage L strain prevalent in Brazil [50]. The animals were monitored daily and euthanized 10 dpi. In those male RM, MAYV RNA was detected in muscle, joint, genital, nervous, cardiovascular, urinary, skin, and lymphoid tissues. Inflammation was observed in small joints, with rashes similar to those in humans [48] and elevated cytokine levels as observed in humans [51], including granulocyte colony stimulating factor (G-CSF), MCP-1, and interleukin (IL)-1 receptor antagonist (RA). Also in 2023, Hamilton *et al*. [11] IV inoculated 12 female cynomolgus macaques with 6 log_10_ PFU MAYV, detecting a 3-4 day. The study used strains from all 3 genetic lineages (N, L, and D [50]). Mean RNA viremias peaked 2 dpi for all lineages and, similar to observations from RM, MAYV RNA was detected in many tissues. Elevated levels of IFN-γ, MCP-1, IL-1RA, IL-10, IL-13, and IL-2 were also detected, similar to human MAYV [51,52]. MAYV lineage D produced significantly higher lymphoid tissue MAYV RNA levels in cynomolgus macaques compared to lineages N and L, while lineage N produced significantly higher RNA levels in the small joints, all at 31 or 33 dpi. Liver and lymph node inflammation were detected, but joint inflammation was not observed 31 or 33 dpi.

Our study builds on this body of research on MAYV in NHP by further defining MAYV viremia magnitude and kinetics, tissue tropism, and histopathologic changes in a larger number of both male and female RM inoculated with a lineage D MAYV strain (IQT4235) not previously tested in NHP. Compared to CHIKV, MAYV viremias in RM are shorter and lower than the 4-6 day average and 5-12 log_10_ PFU/ml peak CHIKV kinetics in macaques (reviewed in [53]). Collectively, the data from this and prior MAYV NHP studies demonstrate that macaques are susceptible to acute MAYV infection, exhibiting short viremias, minor rashes, and mild inflammatory disease in muscles and joints, which mirrors human arthritic symptoms, where MAYV arthralgia occurs commonly in the hand, knee, ankle, foot, and wrist [54]. Alphavirus replication in muscle tissues impacts pathogenesis and persistent disease signs [7,55,56]; for MAYV, this could be influenced by MAYV RNA detection in muscles and other tissues after clearance of infectious virus and viral RNA from blood. The absence of infectious MAYV in tissues 10 or 12 dpi may result from sampling after virus clearance. Isolation of infectious virus from male RM tissues 10 dpi by Weber *et al*. yielded MAYV in several lymph nodes and joints but no other tissues [12]. Detection of MAYV RNA in cerebellum and cerebrospinal fluid in the RM in this study suggests nervous tissue involvement. Neurologic disease signs have not been reported in MAYV patients but occur in CHIKV patients [57,58] and MAYV infects cultured human brain cells [59].

Although prior studies have examined vector competence for *Ae. aegypti* that ingested MAYV from artificial bloodmeals [27–30] and mice [26], this is the first study to expose *Ae. aegypti* to MAYV-infected RM. Even though infectious viremias were detected from 1-3 dpi in all RM and 11 of 12 RM infected at least 1 *Ae. aegypti*, only mosquitoes that fed 2 dpi became infected (caveat: there was 1 mosquito that fed 3 dpi which became infected). These data show there is a narrow temporal window of RM MAYV infectivity for *Ae. aegypti.* Only 2% of all *Ae. aegypti* that bloodfed 2 dpi and 10% of those that developed disseminated infections transmitted infectious MAYV in saliva. We did not observe a correlation between RM viremia 2 dpi and *Ae. aegypti* cohort infection rates. The low transmissibility from RM to *Ae. aegypti* and from *Ae. aegypti* into capillary tubes and the lack of dose response contrasts with patterns from conventional vector competence assessments that use artificial bloodmeals. These studies used mosquitoes from the United States [28,29], Peru [27], or Brazil [30] and bloodmeals with 5-9 log_10_ PFU/ml of MAYV strain IQT4235 which derives from a human in Iquitos, Peru in 2001 (the same strain we used in this study) or TRVL4675 which was isolated from a human in 1954 in Trinidad. Collectively these studies reported higher rates of MAYV transmission than in our study; infectious MAYV was detected in 50-90% of saliva samples collected 7 or 14 dpf of the artificial bloodmeal. The study using Peruvian mosquitoes and IQT4235 [27] also detected a dose-dependent response wherein infection and transmission rates increased with higher bloodmeal titers. These disparities suggest that artificial bloodmeals may overestimate susceptibility to and transmissibility by *Ae. aegypti* for MAYV compared to RM-derived infections, especially at bloodmeal doses that exceed human or RM viremias.

We observed that individual RM were variably infectious to mosquitoes. Intra-host variability in infectiousness is another feature of arbovirus transmission to vectors that artificial bloodmeals may not recapitulate and which has not been evaluated from NHP for any alphavirus including MAYV. The variability was not related to the magnitude of viremia or viral genetic changes in the MAYV genome in RM plasma at the period of mosquito feeding. Given that the animals we used were outbred, we cannot exclude RM genetic factors that condition differential infectiousness of individual animals to mosquitoes. We also did not evaluate cytokine levels or anti-MAYV IgG and IgM titers that may reduce human infectiousness to mosquitoes [60,61]. RM viremia was assessed from blood collected via the femoral vein immediately preceding mosquito presentation on the abdomen of each animal; if levels in the skin and blood-borne virus in the capillary beds in the abdomen differ from the femoral vein, the measured viremia might be different than levels the mosquitoes ingested. Mosquito probing can be both in venules and extra-venular [16]; it is also possible that MAYV in the epidermis was ingested by feeding mosquitoes but would not be represented in viremias reported. We did not sample skin at the time of feeding to assess this possibility. Individual mosquitoes may also have ingested variable bloodmeal volumes, which may have impacted infection success. There are multiple studies for mosquito-borne flaviviruses including dengue (DENV), yellow fever (YFV), and Zika (ZIKV) that show host and virus species both affect infectiousness of an infected vertebrate for a mosquito vector. For DENV, higher viremia is the major determinant of human-to-*Ae. aegypti* transmission [61–64]. This contrasts with our prior NHP studies, where *Ae. albopictus* exposed to experimentally DENV-2-infected cynomolgus macaques and squirrel monkeys were not more likely to become infected after feeding on animals with higher viremias, and, unexpectedly, some aviremic animals even transmitted DENV to *Ae*. *albopictus* [65]. Conversely, we found that rates of infection in *Ae. aegypti* that ingested bloodmeals from YFV-infected cynomolgus macaques [66] and *Ae. albopictus* that ingested bloodmeals from ZIKV-infected squirrel monkeys or cynomolgus macaques [65] correlate with NHP viremia levels.

Our study has several limitations. We used a single strain of MAYV from lineage D and *Ae. aegypti* from a single origin, Los Angeles, California. Only 4 RM per group were used and the sex balance was uneven. Since we exposed the RM to *Ae. aegypti* and mosquito feeding can induce responses in the skin, we cannot exclude mosquito feeding as a cause of skin manifestations. We did not include a group of RM that was sham-inoculated and evaluated histologically; as such, we relied on baseline histopathologic observations in joint and muscle tissues from other RM that were not treated identically to the animals in this study. Our assessments of MAYV RNA levels and tropism in tissues is limited to 10 and 12 dpi, which may miss kinetic changes. We did not present *Ae. aegypti* to RM 1 dpi; this may have overlooked an early period during which the animals were infectious to mosquitoes. The high limit of detection (3.3 and 5.4 log_10_ genomes/ml) for the RT-qPCR assays used for tissues may have resulted in false negatives. The comparison of infectiousness of RM versus artificial bloodmeals used *Ae. aegypti* from two regions; regional variation in vector competence of *Ae. aegypti* from 2 places may bias the comparison.

This study further establishes RM as model of human MAYV and shows that *Ae. aegypti* are capable of transmitting MAYV from infected RM at a very low rate. Poor MAYV transmissibility by *Ae. aegypti* infected from RM may reflect human infectiousness and could, in part, explain why MAYV outbreaks in urban areas are not widespread. Epidemics of CHIKV have been potentiated by adaptation for increased infection and transmission by urban vectors. Mutations in the CHIKV envelope glycoprotein genes have promoted outbreaks since 2005 by increasing infectivity and transmissibility of the virus for *Ae. albopictus* [67,68]. Similarly, a VEEV outbreak in Mexico was likely initiated via envelope gene mutations that augment infectivity for *Ae. (Ochlerotatus) taeniorhynchus* [69]. Given that MAYV is an emerging public health threat [1,18,21], the continued development of NHP models and assessments of transmission to urban *Ae. aegypti* mosquitoes advance our understanding of MAYV disease and the potential for human-*Ae. aegypti*-human cycling.

## ACKNOWLEDGEMENTS

We thank the staff of California National Primate Research Center (CNPRC) Colony Management and Research Services and Clinical Laboratories for their expert technical assistance. We thank Andrew Routh at ClickSeq Technologies for supporting the sequencing aspects of this project. This study was funded by a CNPRC Pilot Program Award (to SLR); Award P51OD011107 from the Office of Research Infrastructure Program, Office of The Director, National Institutes of Health to the California National Primate Research Center, and by the World Reference Center for Emerging Viruses and Arboviruses, NIH grant R24AI120942. AJM was supported by the National Institutes of Health, National Institute of Allergy and Infectious Diseases Animal Models of Infectious Disease T32AI060555 as well as the University of California Davis School of Veterinary Medicine Graduate Student Support Program Fellowship.

## AUTHOR CONTRIBUTIONS

Conceptualization: AJM, KKVR, WL, JKW, RKC, SCW, SLR, LLC

Data Curation: AJM, KKVR, WL, JKW, JU, PNC, KJO, CSM

Formal Analysis: AJM, LLC

Funding Acquisition: SLR, LLC, KKVR

Investigation: AJM, KKVR, WL, SA, RL, JU, JKW, PNC, KJO, CSM

Methodology: AJM

Project Administration: KKVR, SLR, LLC Resources: SCW

Supervision: KKVR, SLR, LLC Validation: AJM, KKVR, LLC

Visualization: AJM, LLC

Writing-Original Draft Preparation: AJM, LLC

Writing-Review and Editing: AJM, KKVR, WL, SA, RL, JU, PNC, KJO, CSM, RKC, SW, SLR, LLC

